# TGFβ receptor inhibition unleashes interferon-β production by tumor-associated macrophages and enhances radiotherapy efficacy

**DOI:** 10.1101/2022.01.17.476557

**Authors:** Pauline Hamon, Marine Gerbé de Thoré, Marion Classe, Nicolas Signolle, Winchygn Liu, Olivia Bawa, Lydia Meziani, Céline Clémenson, Fabien Milliat, Eric Deutsch, Michele Mondini

## Abstract

**Background:** Transforming growth factor-beta (TGFβ) can limit the efficacy of cancer treatments, including radiotherapy (RT), by inducing an immunosuppressive tumor environment. The association of TGFβ with impaired T cell infiltration and antitumor immunity is known, but the mechanisms by which TGFβ participates in immune cell exclusion and limits the efficacy of antitumor therapies warrant further investigations.

**Methods:** We used the clinically relevant TGFβ receptor 2 (TGFβR2)-neutralizing antibody MT1 and the small molecule TGFβR1 inhibitor LY3200882 and evaluated their efficacy in combination with RT against murine orthotopic models of head and neck and lung cancer.

**Results:** We demonstrated that TGFβ pathway inhibition strongly increased the efficacy of RT. TGFβR2 antibody upregulated interferon beta (IFNβ) expression in tumor-associated macrophages (TAMs) within the irradiated tumors and favored T cell infiltration at the periphery and within the core of the tumor lesions. We highlighted that both the antitumor efficacy and inhibition of immune exclusion observed with the combination of MT1 and RT were dependent on type I interferon signaling.

**Conclusions:** These data shed new light on the role of TGFβ in limiting the efficacy of RT, identifying a novel mechanism involving the inhibition of macrophage-derived type I interferon production, and fostering the use of TGFβR inhibition in combination with RT in therapeutic strategies for the management of head and neck and lung cancer.

## BACKGROUND

Radiotherapy (RT) contributes to the treatment of more than 60% of all cancer patients, including a significant proportion of head and neck squamous cell carcinoma (HNSCC) and lung cancer patients. It is widely accepted that RT efficacy relies not only on its cytotoxic activity but also on its widespread effects on the tumor microenvironment and notably on the modulation of the immune response. In the last decade, a large amount of preclinical data have been generated, demonstrating that RT can trigger both adaptive and innate immune responses toward antitumor activity. Indeed, RT can induce so-called “immunogenic cell death” (ICD), a cell death modality that fosters a T-cell mediated immune response against tumor antigens [1]. RT can also activate the STING cytosolic DNA-sensing pathway, triggering the interferon (IFN) response and thus stimulating an antitumor adaptive immune response, which plays a pivotal role in mediating the antitumor effects of RT [2]. On the other hand, it has also been shown that RT can trigger immunosuppressive signals. For instance, we and others have shown that upon irradiation, an influx of monocytes and regulatory T cells (Tregs) into the tumor, which are recruited via the CCL2/CCR2 pathway, restrain the antitumor efficacy of RT [3,4]. Several other cellular and soluble factors contributing to the development of an immunosuppressive environment can limit the efficacy of cancer treatments. Among these, transforming growth factor-beta (TGFβ), a pleiotropic cytokine involved in numerous pathways of tumor growth, appears to play a prominent role. TGFβ can promote tumor metastasis and progression by increasing the motility of cancer cells, facilitating the process of epithelial-to-mesenchymal transition (EMT), inducing angiogenesis and recruiting myeloid cells within the tumor microenvironment (TME) [5]. TGFβ also exerts immunosuppressive effects through the induction of Tregs and the inhibition of cytotoxic CD8 T cell (CTL) activities [6]. There is also evidence that TGFβ signaling limits T cell infiltration in the TME and the response to immune checkpoint blockade [7,8]. A recent report demonstrated that CD8 T cell trafficking to the tumor is limited by TGFβ through inhibition of CXCR3 expression in CD8^+^ T cells [9]. As TGFβ is implicated in several different pathways, the mechanisms by which TGFβ participates in immune cell exclusion and limits the efficacy of antitumor therapies warrant further investigations.

TGFβ is secreted as a latent complex in the extracellular matrix until external stimuli dissociate it from latency-associated peptide (LAP) [10]. RT-generated reactive oxygen species (ROS) modify the LAP-TGFβ complex to release active cytokines [11]. The increased bioavailability of TGFβ within the TME of irradiated tumors can contribute to the generation of an immunosuppressive environment that hampers the efficacy of RT. Accordingly, antibody-mediated TGFβ neutralization synergizes with RT and generates CD8 T cell responses to multiple endogenous tumor antigens in poorly immunogenic mouse carcinomas [12].

Ligand interactions with transforming growth factor β receptor type I (TGFβR1) and type II (TGFβR2) complexes are responsible for the biological activity of TGFβ. Binding of TGFβ ligands (TGFβ 1, 2, or 3) to TGFβR2 induces phosphorylation of the receptor’s serine/threonine kinase domain, and then the ligand-receptor forms a heterotrimeric phosphoprotein complex with TGFβR1, triggering the activation of TGFβ signaling pathways [13]. Inhibition of TGFβR1 or R2 impairs the SMAD-dependent signal transduction elicited by active TGFβ [14]. Targeting TGFβR2 with a specific neutralizing antibody has shown significant efficacy against primary tumor growth and metastasis [15,16].

Here, we identified a novel mechanism for TGFβ-mediated immune suppression after RT. Such mechanism involves the inhibition of type I interferon production by myeloid cells. Using the clinically relevant TGFβR2 neutralizing antibody MT1 (the murine equivalent of the human LY3022859 antibody [17]) and the small molecule TGFβR1 inhibitor (LY3200882 [18]), we demonstrated that TGFβ pathway inhibition strongly increased the efficacy of RT against two different murine orthotopic models of head and neck carcinoma, as well as an orthotopic model of lung cancer. TGFβR2 antibody unleashed IFNβ synthesis by macrophages and significantly favored T cell infiltration not only at the tumor periphery but also throughout all tumor regions, including the core. Finally, we demonstrated that both the antitumor efficacy and the inhibition of immune exclusion observed with the combination of anti-TGFβR2 and RT were dependent on type I interferon signaling. These data shed new light on the crosstalk between the TGFβ and type I IFN pathways and foster the use of TGFβR inhibition in combination with RT in therapeutic strategies for the management of head and neck and lung cancer.

## METHODS

### Cell lines

The TC-1 tumor cell line was derived from primary lung epithelial cells of a C57Bl6 mouse cotransformed with HPV-16 oncoproteins E6 and E7 and the c-Ha-ras oncogene [19]. Firefly luciferase-expressing TC1 (TC1/Luc) cells were kindly provided by T.C. Wu (Johns Hopkins Medical Institutions, Baltimore, MD, USA) in 2009. The RAW 264.7 monocyte/macrophage cell line was obtained from ATCC (Manassas, VA, USA). The head and neck murine SCC VII cells were kindly provided by C. Brunner (Universitätsklinik Ulm, Ulm, Germany). Lewis lung carcinoma cells expressing luciferase (LL2/Luc) were purchased from Caliper Life Sciences (Hopkinton, MA, USA). Cell lines were routinely screened for mycoplasma contaminations using the MycoAlert mycoplasma detection kit (Lonza).

### Animal models

Seven- to eight-week-old C57BL/6 and C3H/HeN female mice were purchased from Janvier CERT (Le Genest St. Isle, France), and athymic *nude-nu/nu* mice were bred at the Gustave Roussy animal facility (Plateforme d’Evaluation Preclinique, PFEP). All animals were housed at PFEP and included in experiments after at least one week of acclimatization period. To establish a head and neck tumor models, syngeneic tumor grafts were initiated by injection of 50 μL of PBS suspension containing 5×10^5^ TC1/Luc or SCC VII cells at submucosal sites on the right inner lips of C57Bl/6mice or C3H/HeN mice respectively, as previously described [20]. For orthotopic lung tumors, the skin was incised, 100,000 LL2/Luc cells (in PBS + Matrigel (BD Biosciences), 10 μl) were injected directly into the lung through the pleura and then the wound was closed by suture clips [21]. Throughout the study, researchers were aware of the group allocation at any stage of the experiment or data analysis. The health, weight and behavior of the mice were assessed three times per week. Mice were euthanized upon the presentation of defined criteria (tumor size and bioluminescent signal, loss of >20% of the initial weight), and a survival time was recorded to perform a survival analysis for the treatment groups.

### Irradiation

For head and neck tumors, RT-treated mice received single-beam local irradiation of the head and neck region 7 days (TC1/Luc model) or 9 days (SCC VII model) after tumor inoculation using a 200 kV Varian X-ray irradiator. Mice bearing lung tumors received a whole thorax irradiation. Selective irradiation was performed by the interposition of a 4-cm-thick lead shield on a schedule delivering 8 Gy (for TC1/Luc) or 12 Gy (for SCC VII and LL2/Luc) in a single fraction at a dose rate of 1.08 Gy/min.

### Antibodies and treatments

The anti-TGFβR2 antibody MT1, and the rat IgG2a isotype control were supplied by Eli Lilly, New York, NY, USA. Antibodies were administered intraperitoneally (i.p.) at 40 mg/kg for MT1 once a week starting at the indicated time point. Anti-interferon-α/β receptor (IFNAR) antibody (clone MAR1-5A3) and mouse IgG1 isotype control (clone MOPC-21) were purchased from BioXcell and administered i.p. at 200 μg/mouse before RT and then three times per week for a total of four injections. Mice received i.p. injections of 100 μg per mouse of anti-CD8 mAb (clone 2.43; BioXCell) one day before RT to deplete CD8 T cells. The TGFβRI inhibitor LY3200882 was supplied by Eli Lilly. This inhibitor was reconstituted in hydroxyethyl-cellulose (HEC) 0.25% Tween 80 and administered by oral gavage at 75 mg/kg twice a day starting at D7 for 2 weeks. HEC 0.25% Tween 80 as a vehicle was administered at the same frequency to the control groups.

To deplete tumor macrophages, mice bearing TC1/Luc head and neck tumors received intratumor injection of 20 μl of clodronate liposomes (clodrosomes, Liposoma BV) or PBS-loaded liposomes (Liposoma BV), used as control, twenty minutes before irradiation, as previously described [22].

### Histological analysis

Tumors were fixed in 4% buffered paraformaldehyde (PFA), paraffin embedded, and then cut into 4 μm sections. Immunostaining was performed using a Ventana Benchmark^®^ automaton (Ventana, AZ, USA) using anti-CD8 (Cell Signaling, 98941) anti-CD31 (Abcam, ab28364) and anti-CAIX (Abcam, ab15086) antibodies, and digitized using a slide scanner (Olympus VS120, Olympus, Japan). Quantification of stained cells was performed by image analysis using Qupath software. A semiquantitative analysis of CD8^+^ infiltration was performed by a head and neck pathologist at the tumor invasive margin and within the tumor core. CD8^+^ infiltrate was scored from 0 to 3 (score 0: none or negligible even at 20x magnification; score 1: scarce infiltrate observed only at a moderate magnification (10x); score 2: moderate observed at a low magnification (5x); score 3: abundant infiltrate easily observed at low magnification (5x)).

Tumor regression (percent of viable tumor cells) was evaluated by a head and neck pathologist according to the usual criteria used in daily practice conditions [23].

### *In vivo* imaging

To monitor tumor growth, bioluminescence imaging was performed on the indicated days using the Xenogen *In Vivo* Imaging System 50 (IVIS; Perkin Elmer) as previously described and analyzed using Living Image analysis software (Perkin Elmer) [20]. Briefly, mice were injected with D-luciferin i.p. (150 mg/kg) and anesthetized with isoflurane 10 min before imaging.

### Flow cytometry cell analysis

Phenotypic characterization of murine cells was performed using an LSR Fortessa instrument (Becton Dickinson) with DIVA^®^ Flow Cytometry software. For data analysis, FlowJo software (Tree Star Inc.) was used. Five days post radiotherapy, blood was drawn in the cheek with a lithium-heparin minivette POCT (Sarstedt, Germany) and directly stained with antibodies. After staining, erythrocytes were lysed in lysis buffer (BD Pharm Lyse^™^, BD Biosciences) and resuspended in FACS buffer (Stain Buffer FBS, BD Biosciences). At the same time, TC1/Luc oral tumors were digested in RPMI medium (Gibco, Invitrogen, France) supplemented with collagenase IV (1 mg/mL, Sigma) and DNase 1 (1 mg/ml) for 30 min at 37 °C, resuspended in PBS and filtered using a 40 μm-pore cell strainer (BD Biosciences). After centrifugation at 1500 rpm for 5 min at 4 °C, the cells were filtered again using a 70 μm-pore cell strainer (BD Biosciences) in PBS. Surface staining was performed by incubating 1/10^th^ of the cell suspension with 1 μg/mL purified anti-CD16/32 (clone 2.4G2, BD Biosciences) for 10 min at 4 °C followed by an additional 20 min incubation with appropriate dilution of the surface marker antibodies. Cells were then washed once in FACS buffer and directly analyzed by flow cytometry. The panels of antibodies used were: anti-CD11b (clone M1/70), anti-Ly6C (clone AL-21), anti-Siglec-F (clone E50-2440), anti-CD8 (clone 53-6.7), anti-CD25 (clone PC61), anti-IFNγ, anti-PD-L1 (clone MIH5), anti-Foxp3 (clone MF23), all from BD Biosciences; anti-Ly6G (clone 1A8), anti-NK1.1 (clone PK136), anti-I-A/I-E (clone 2G9), anti-CD11c (clone N418), anti-CD45 (clone 30-F11), anti-CD4 (clone RM4-5), anti-CD64 (clone X54-5/7.1) all from BioLegend. For cytokine staining, cells were pre-incubated for 2 h with the Cell Activation Cocktail containing Brefeldin A according to the manufacturer’s instructions (BioLegend). After surface staining, the cells were fixed in 4% PFA for 15 min, washed in Perm/Wash solution (BD Biosciences) and incubated for 30 min at room temperature (RT) in the presence of anti-IFNγ. For intracellular staining without preactivation, the Foxp3/Transcription Factor Staining Buffer Set (BD Biosciences), anti-Foxp3 and anti-CTLA-4 were used according to the manufacturer’s instructions. Samples were washed in FACS buffer and resuspended in 200 μL FACS buffer before acquisition.

Calculation of absolute cell number was performed by adding to each vial a fixed number (10,000) of nonfluorescent 10 μm polybead^®^ carboxylate microspheres (Polysciences, IL, USA) according to the formula: Nb of cells = (Nb of acquired cells x 10,000)/(Nb of acquired beads). The number of cells obtained for each sample was normalized per mg of tissue.

### Flow cytometry cell sorting

Tumor-associated macrophages were sorted using the ARIA Fusion-UV cell sorter (BD Biosciences). TC1/Luc oral tumors were irradiated and treated with MT1 or IgG 8 days after cell inoculation. Tumors were collected 1 day after irradiation. Tumors were digested in RPMI medium (Gibco, Invitrogen, France) supplemented with collagenase IV (1 mg/mL, Sigma) and DNase 1 (1 mg/ml) for 30 min at 37 °C, entirely resuspended in PBS and filtered using a 40 μm-pore cell strainer (BD Biosciences). After centrifugation at 1500 rpm for 5 min at 4 °C, the cells were filtered again using a 70 μm-pore cell strainer (BD Biosciences) in PBS. Surface staining was performed by incubating the cell suspension with 1 μg/mL purified anti-CD16/32 (2.4G2, BD Biosciences) for 10 min at 4 °C followed by an additional 20 min incubation with appropriate dilution of the surface marker antibodies. Cells were then washed once in PBS and directly sorted by ARIA Fusion-UV. TAMs were stained with anti-CD11b (clone M1/70), anti-Ly6C (clone AL-21), anti-CD64 (clone X54-5/7.1) and anti-Ly6G (clone 1A8) and sorted as CD11b^+^ CD64^+^ Ly6C^low^ Ly6G^low^ with a purity >95%.

### Bone marrow–derived macrophages cultures

The femur and tibia of C57Bl/6 mice were flushed, and bone marrow cells was obtained using a previously described protocol [24]. The adherent cells were incubated in medium supplemented with recombinant mouse M-CSF (R&D) at a concentration of 250 ng/mL for five days to induce anti-inflammatory bone marrow–derived macrophages (BMDMs) as previously described [25].

### Cytokine/chemokine array

The cytokine and chemokine concentrations in tumor tissues were profiled using the Mouse Focused 10-Plex (MDF10) at Eve Technologies Corporation (AB, Canada). Protein extracts from tumor tissues were prepared by homogenization in RIPA buffer (Sigma-Aldrich, USA) containing a protease inhibitor cocktail (Roche, Switzerland) using Biomasher disposable homogenizers (Nippi, Japan). Protein extracts were diluted to 4 μg/μL, and the multiplex immunoassay was analyzed with the BioPlex 200 instrument (Bio-Rad, USA). Cytokine and chemokine concentrations were calculated based on the standard curve generated using the standards included in the kit.

### Elisa

RAW 264.7 cells or BMDMs were cultured as described above and then treated with recombinant mouse TGFβ1 (Biolegend). Fifteen minutes later, the cells were irradiated with 8 Gy and then treated with MT1 or IgG at 0.1 μg/mL. Treatments with TGFβ and MT1 or IgG were renewed 24 h after irradiation. Supernatants were collected one day later. Supernatants of BMDMs were 5-fold concentrated using the Amicon Ultra-0,5 Membrane Ultracel-10 columns (Merck Millipore) and the IFNβ concentration was evaluated using the VeriKine Mouse Interferon Beta HS ELISA Kit (PBL assay science).

### Quantitative PCR

Total RNA was extracted from sorted macrophages using the Qiagen Micro Plus RNeasy kit. RNA was retrotranscribed using SuperScript VILO Mix (Invitrogen). qRT-PCR analysis of IFNβ was performed using Fast Advanced TaqMan Master Mix (Invitrogen) with the predesigned TaqMan assays Mm00439552_s1 (mIFNβ) and Mm00437762_m1 (beta2 microglobulin, B2M) as housekeeping genes (Invitrogen) using the 7500 RealTime PCR System (Life Technologies).

### Statistical analyses

Statistical analyses were performed using Prism Version 9 (GraphPad, CA, USA). Survival data were analyzed using the Kaplan-Meier and log-rank tests for survival distribution. The bioluminescent signal (serving as a measurement of tumor size), immune infiltrate and cytokine levels were analyzed using appropriate tests as indicated. Multigroup analyses of variances were performed using one-way ANOVA followed by indicated post-tests or the Kruskal-Wallis test followed by Dunn’s multiple comparisons test. For simple comparison analysis, the unpaired Student’s t-test with Welch’s correction was used to compare parametric distributions, while non-parametric testing was performed using the Mann-Whitney test. *: p<0.05; **: p<0.01; ***: p<0.001; ****: p<0.0001; ns: nonsignificant.

## RESULTS

### TGFβ receptor blockade increases radiotherapy efficacy in head and neck and lung tumors

We previously demonstrated that 7.5 Gy irradiation exerted a limited effect on murine oral tumors when obtained by orthotopic injection of the TC1/Luc cell line at a submucosal site of the inner lip in syngeneic mice [4,20]. Using the same model, we showed that two weekly administrations of the anti-TGFβR2 antibody (MT1) were sufficient to significantly improve the antitumor efficacy of RT (Fig. 1A-B). Bioluminescent *in vivo* imaging showed that MT1 and RT as monotherapies slightly reduced tumor growth, while the combined treatment resulted in shrinkage of most of the tumors with long-lasting complete responses (Fig. 1B). Accordingly, mouse survival was significantly increased in the group receiving the RT and MT1 combination, with a median survival of 36 days, compared to 11.5 days and 17 days for the IgG-treated and RT groups, respectively (Fig. 1A). We further confirmed that TGFβR2 blockade improves the efficacy of RT in another orthotopic model of head and neck cancer obtained by the intramucosal injection of the HPV-negative SCC VII cells. The combination of MT1 treatment with a 12Gy local irradiation significantly increased the mouse survival when compared to RT alone (Fig. 1C). The antitumor activity of MT1 in combination with RT was not restricted to the oral tumor setting, as the survival of mice bearing LL2/Luc lung orthotopic tumors was improved when they were treated with the RT and MT1 combination (Fig. 1D). Finally, the inhibition of TGFβR1 using the small molecule LY3200882 confirmed that targeting the TGFβ pathway increases the efficacy of RT, even if the effects were lower than those observed with MT1 in the TC1/Luc model (Supp. Fig. 1A-C).

**Figure 1:**
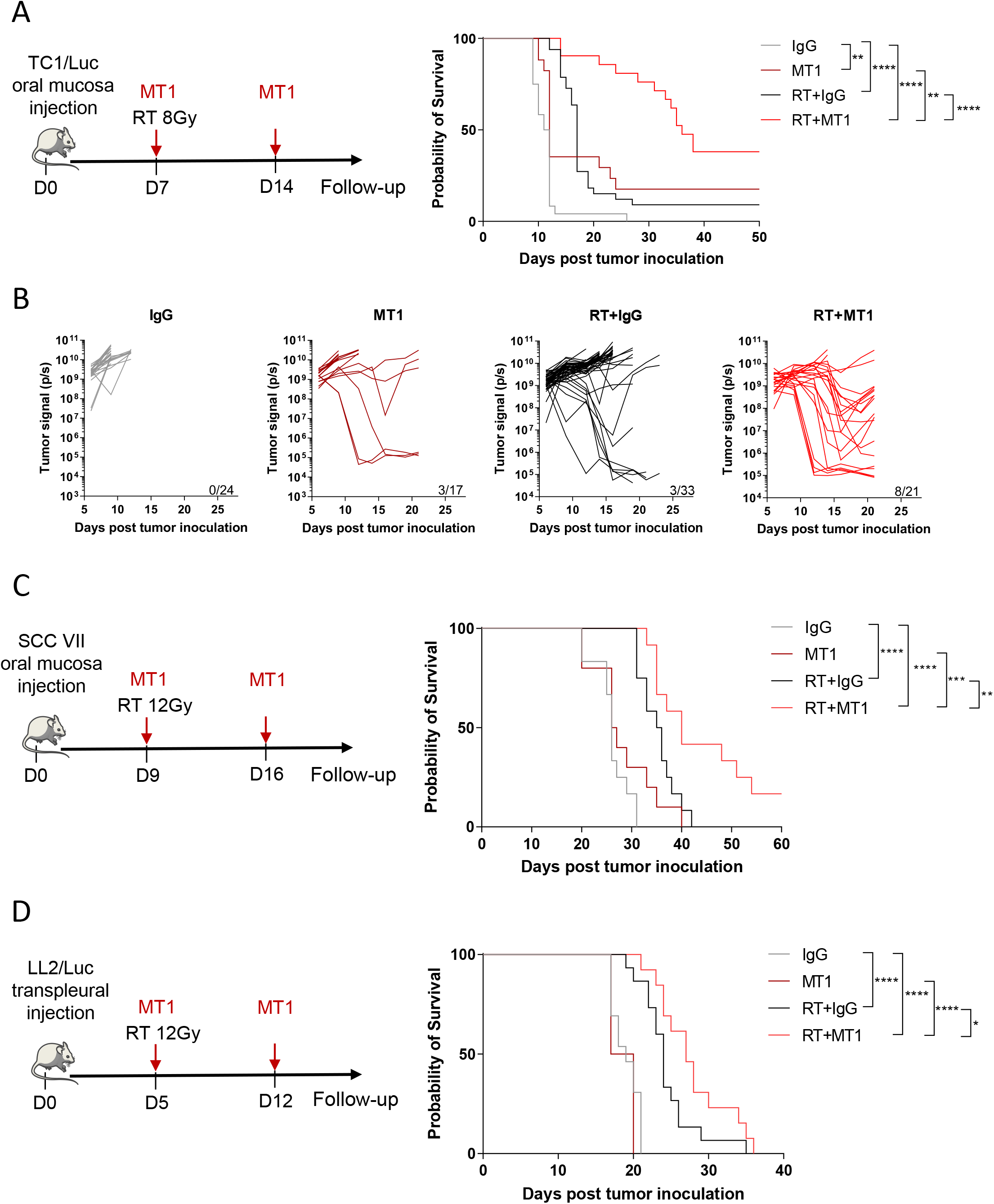
TGFβ receptor blockade increases radiotherapy efficacy in head and neck and lung tumors. (**A**) Left panel shows the treatment scheme used for theTC1/Luc head and neck model, with irradiation at 8 Gy at day 7 and two weekly injection of MT1. Right panel presents the Kaplan-Meier survival curves of the different treatment groups. (**B**) *In vivo* bioluminescent imaging was performed to follow tumor growth, and quantification of the bioluminescent signal from individual mice was performed at different time points post treatment (For A and B: n = 17-33 mice/group from 4 independent experiments). (**C**) Left panel presents the treatment scheme used for the SCC VII head and neck mouse model, with irradiation at 12 Gy at day 9 and two weekly injection of MT1. Right panel shows the Kaplan-Meier survival curves (n= 10-12 mice/group from 3 independent experiments). (**D**) Left panel shows the treatment scheme used for the LL2/Luc lung orthotopic mouse model and includes the irradiation at 12 Gy at day 5 and two weekly injection of MT1. Right panel presents the Kaplan-Meier survival curves (n= 4-15 mice/group from 2 independent experiments). For all panels: *: p<0.05; **: p<0.01; ***: p<0.001; ****: p<0.0001 (A; C-D, log-rank test,).

Altogether, these data showed that the combination of radiotherapy with TGFBR inhibition is beneficial in multiple cancer type as head and neck and lung tumors.

### TGFβR2 inhibition by MT1 fosters an antitumor immune environment in irradiated tumors

Local radiotherapy has been described to induce immunogenic cell death and the recruitment of immune cells to the tumor site [1]. As TGFβ is immunosuppressive, we hypothesized that the combination of MT1 would synergize with RT and modify T cell infiltration and function. We analyzed the immune environment of TC1/Luc oral tumors 6 days after treatment administration. Irradiation slightly upregulated the number of tumor-infiltrating CD8^+^ T cells compared to control mice, while it was strongly enhanced by the combination with MT1 antibody, reaching nearly 8-fold the level observed in untreated tumors (Fig. 2A). Similar recruitment was observed for CD4^+^ conventional T cells (Tconv), Tregs and NK cells in the RT+MT1 group (Fig. 2A). In the irradiated groups, tumor-infiltrating CD8^+^ T cells were activated, as demonstrated by an increase in the proportion of IFNγ-producing cells (Fig. 2B). Accordingly, cytokine profiling of tumor protein extracts showed an increased amount of IFNγ in the tumor environment in mice treated with the RT and MT1 combination and a trend for other proinflammatory cytokines, such as IL-1β and IL-6, as well as TNFα, while IL-4 and IL-12 were unaffected by the treatment (Fig. 2C).

**Figure 2:**
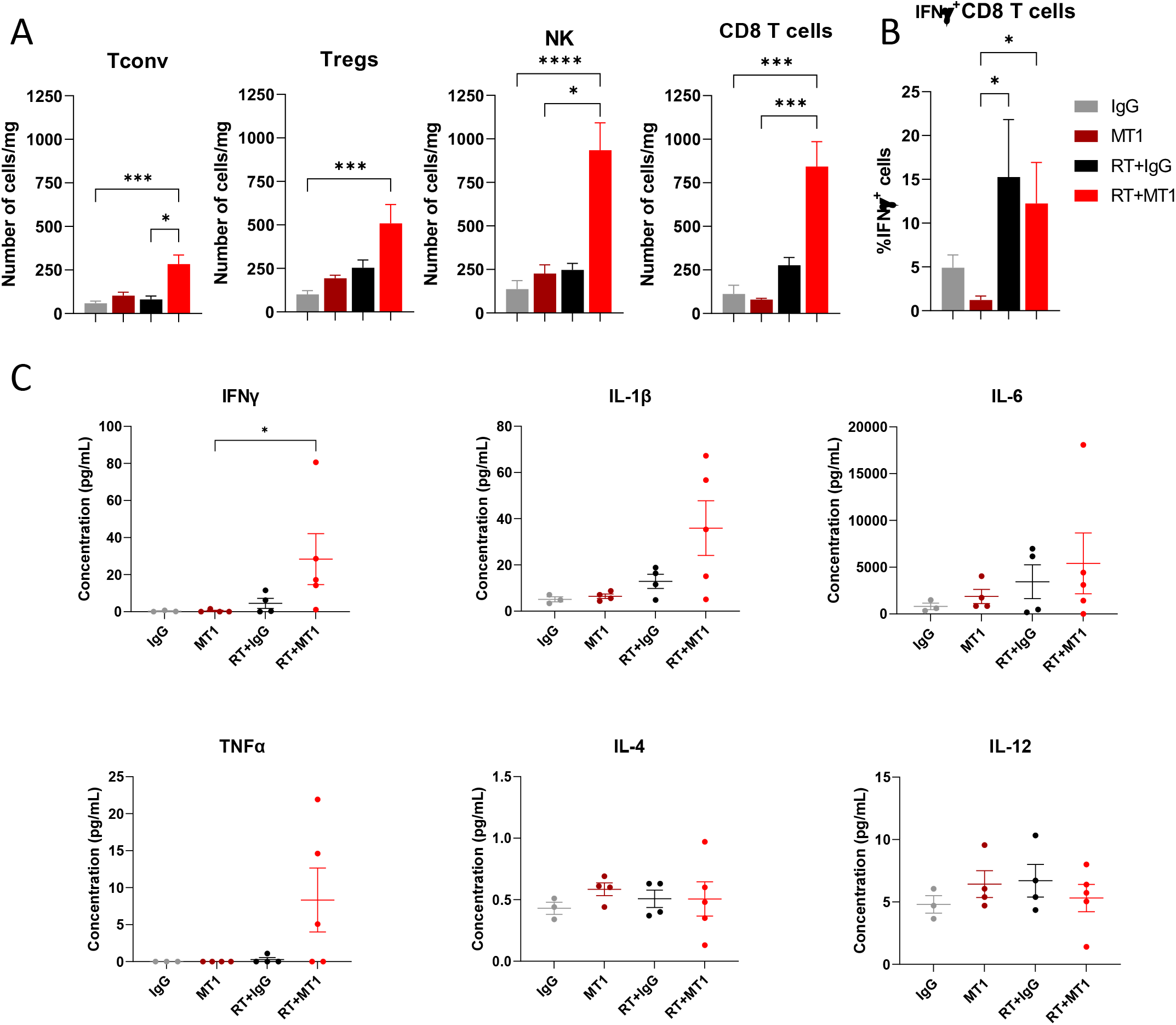
TGFβR2 inhibition by MT1 fosters an antitumor immune environment in irradiated tumors. Mice bearing TC1/Luc tumors were treated as in Fig. 1A, and tumor specimens were collected 6 days post RT to perform cytofluorimetry analyses. (**A**) Histograms showing the number of cells/mg in the tumor for conventional CD4^+^ T cells (Tconv), regulatory T cells (Tregs), natural killer cells (NK) and CD8^+^ T cells. (**B**) Percent of CD8^+^ T cells expressing IFNγ in the different groups (for C and D: n = 8-9 mice from 3 independent experiments, mean ± SEM is reported). (**C**) Cytokine profiling of whole-tumor protein extracts performed 6 days after radiotherapy. For all panels: *: p<0.05; ***: p<0.001; ****: p<0.0001 (Kruskal-Wallis with Dunn’s multiple comparison test).

### MT1 favors in-depth cytotoxic T-cell infiltration of irradiated tumors

A strong limitation of T cell-mediated antitumor function is the quality of recruitment and infiltration of cytotoxic CD8 T cells within the tumor. “Cold” HNSCCs with poor lymphoid infiltration have the worst overall survival, including when treated with RT [26]. Histological examinations of TC1/Luc tumor tissues 6 days post treatment confirmed that the combination of RT and MT1 exerted a significant antitumor effect, with a 70% reduction of the remaining viable tumor tissue compared to IgG control (Fig. 3A), which is in agreement with data observed by *in vivo* bioluminescent imaging (Fig. 1B). MT1 treatment induced a trend in increasing the vascularization of the tumors, as shown by immunohistochemistry (IHC) CD31 staining (Supp. Fig. 2A-B), as well as a reduction of tumor hypoxia, as shown by the reduction of the extent of staining of the hypoxic marker CAIX (Supp. Fig. 2C-D). IHC staining for CD8 showed an accumulation of CD8^+^ T cells in the combination group (see representative images in Fig. 3B and quantification in Fig. 3C), confirming the data obtained by flow cytometry (Fig. 2A). Interestingly, while the density of CD8^+^ T cells increased at the periphery of the tumor in all treatment groups when compared to control mice (Fig. 3D), the combination of RT and MT1 induced a higher accumulation in the tumor core than the single treatments (Fig. 3E). Automated quantification of the spatial distribution of CD8^+^ T cells showed that their density steadily decreased from the periphery to the core in the control and monotherapy groups but remained stable throughout all RT+MT1-treated tumors (Fig. 3F, left graph). Therefore, the ratio of CD8^+^ T cell density in the RT+MT1 *vs* IgG group strongly increased when going toward the inner tumor core (Fig. 3F, right graph). This demonstrates that the combination treatment with MT1 favors the accumulation of CD8^+^ T cells even within poorly infiltrated tumor regions.

**Figure 3:**
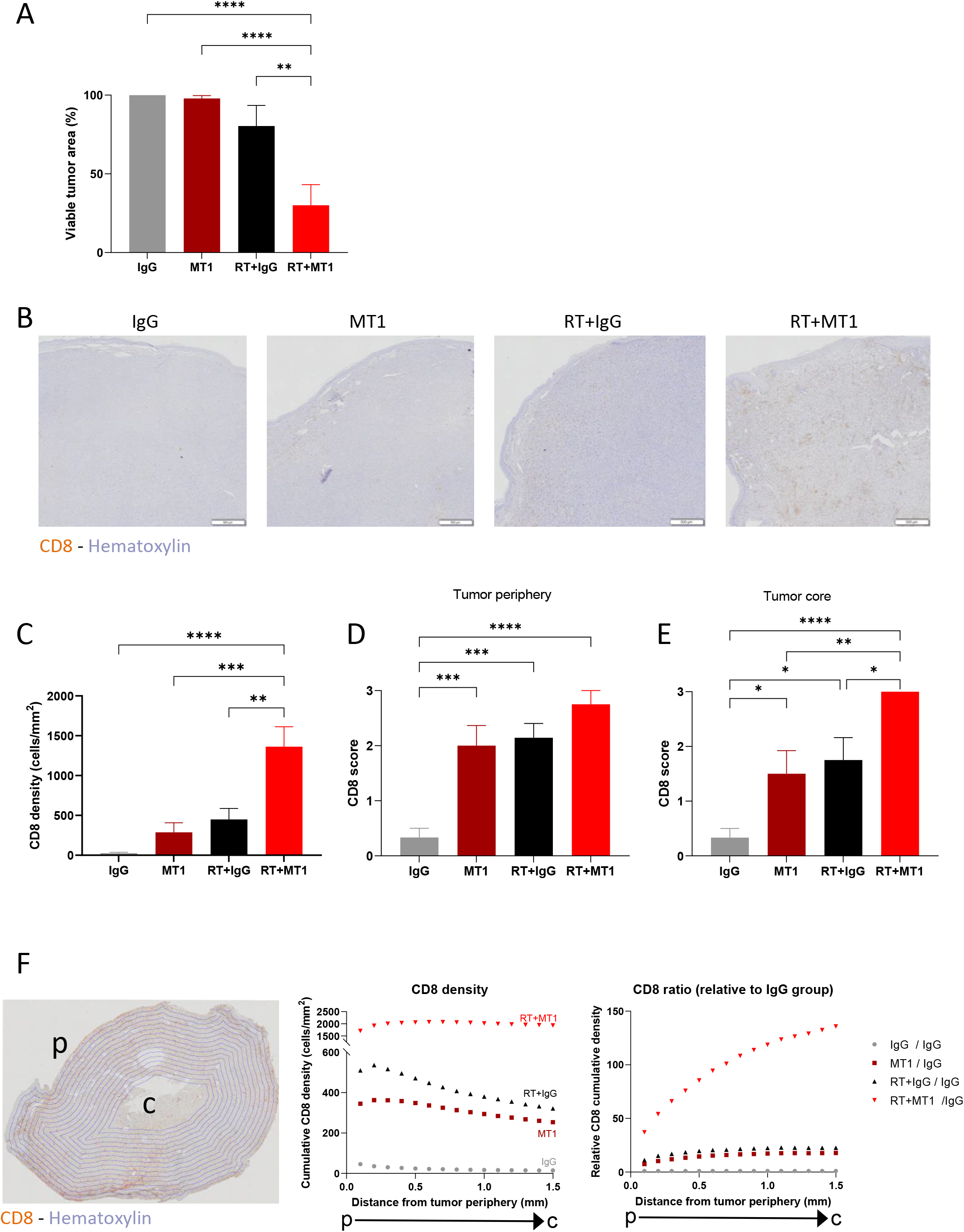
MT1 favors in-depth cytotoxic T-cell infiltration in irradiated tumors. Mice bearing TC1/Luc tumors were treated as in Fig. 1A, and tumor specimens were collected 6 days post RT to perform histological analyses. (**A**) Histogram representing the percent of viable tumor for each group quantified by histological examination. (**B**) Representative images of CD8 staining by immunohistochemistry on head and neck tumors. (**C**) Quantification of CD8 T cell density per mm^2^. (**D-E**) Semiquantitative scoring of the CD8 density at the tumor periphery (D) and at the tumor core (E). (**F**) Scans of tumor sections were segmented in 0.1 mm concentric layers, as shown in the representative image (left panel). Quantification of CD8 T cell density was performed in each layer, and cumulative densities starting from the tumor periphery (p) to the tumor core (c) for each group are reported in the middle panel. The right panel shows the ratio of CD8 in each group relative to the IgG group. For all panels, n = 8-9 mice from 2 independent experiments, *: p<0.05; **: p<0.01; ***: p<0.001; ****: p<0.0001 (one-way ANOVA with Tukey’s multiple comparison test).

### MT1 and RT combination efficacy depends on CD8^+^ T cells

To validate that the efficacy of the combination of MT1 and RT relied on increased activation of the antitumor immune response, we injected TC1/Luc cells into the oral mucosa of nude mice. Analysis of tumor growth and survival (Supp. Fig. 3A-B) showed that MT1 treatment had no effects in immunodeficient mice, without any improvement in the efficacy of radiotherapy. We next demonstrated that the activity of tumor-infiltrating CD8^+^ T cells was required to mediate the effects of TGFβR2 inhibition. Indeed, when we depleted this population using an anti-CD8 antibodies, the efficacy of the combined treatment was completely lost, and none of the mice responded to the combination of RT with MT1 (Fig. 4A). Accordingly, the survival of the mice was significantly reduced when CD8 antibodies were administered to mice treated with RT and MT1 combination, reaching a level similar to that observed in mice treated with RT alone (Fig. 4B). We next evaluated if the activation of the immune system by the combination of RT and MT1 could result in the induction of long-term antitumor immune memory. To test this hypothesis, 10 mice treated with RT and MT1 that underwent complete tumor clearance and that were still tumor free 60 days after tumor engrafting, were challenged again with TC1/Luc cells. None of the mice developed a tumor, while all treatment-naïve mice (used as positive controls) displayed a tumor growth as expected (Fig. 4C).

**Figure 4:**
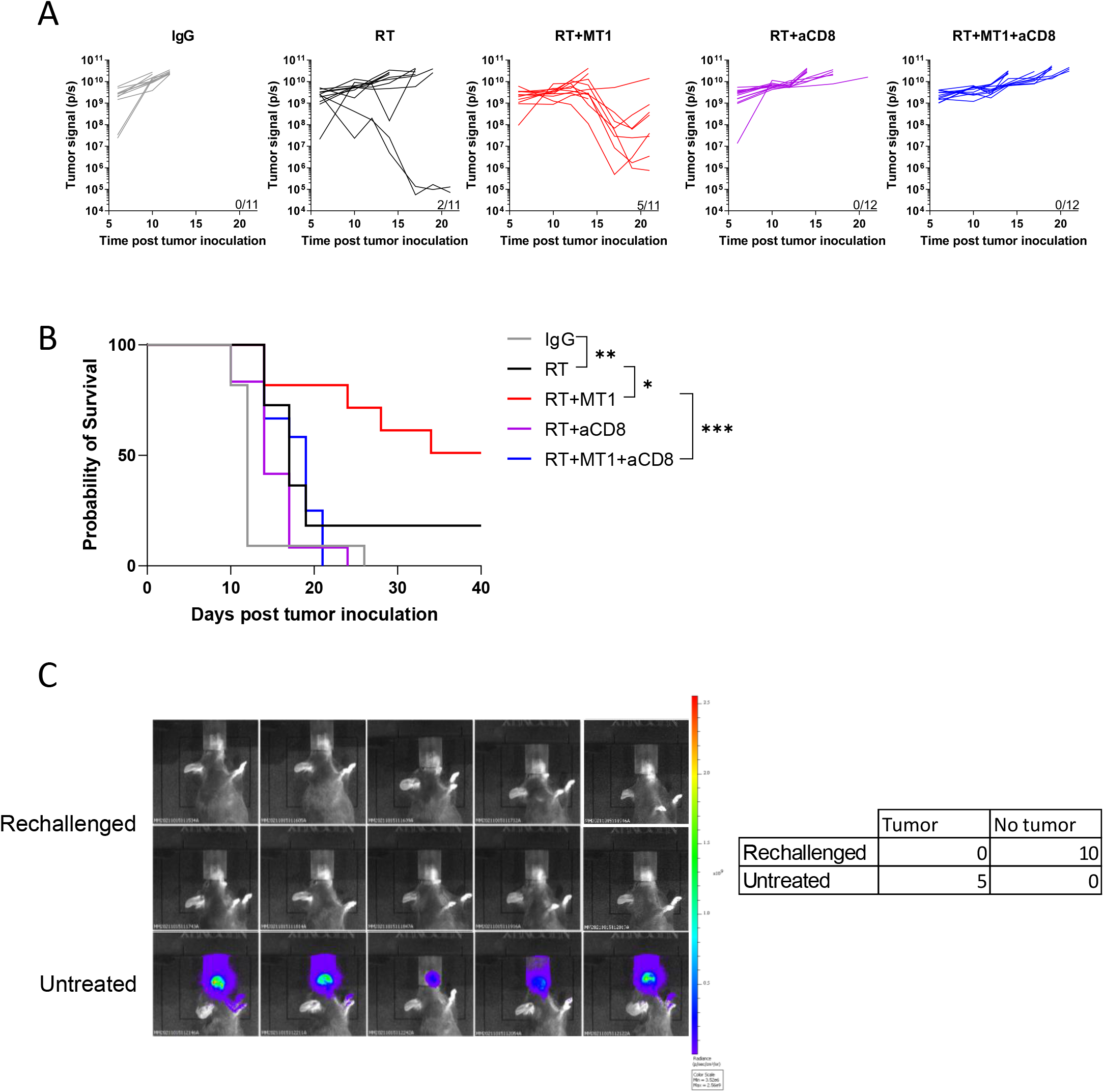
MT1 and RT combination relies on CD8^+^ T cells, and induces long-term antitumor memory. (**A**) Quantification of the bioluminescent signal from individual TC1/luc tumors was performed at different time points post treatment in the different groups treated with or without anti-CD8 antibody (aCD8). (**B**) Kaplan-Meier mouse survival curves (For A-B, n = 11-12 mice/group from 2 independent experiments). (**C**) Left panel shows images of the bioluminescent signal 10 days after TC1/Luc injection in 10 mice previously treated with RT and MT1 that underwent complete TC1/Luc tumor clearance and that were still tumor free 60 days after tumor grafting (Rechallenged), and in 5 treatment-naïve mice (Untreated). Right panel present the summary table of mice that developed or not tumors. For all panels: *: p<0.05; **: p<0.01; ***: p<0.001 (log-rank test).

### TGFβR2 inhibition triggers the production of IFNβ by irradiated macrophages

Together with increased T cell infiltration, RT slightly increased the number of tumor-associated macrophages (TAMs) 6 days after radiotherapy in TC1/Luc tumors (Fig. 5A), in agreement with our previous report [4], which was not further modulated by treatment with MT1. A trend of an increase in Ly6C^high^ monocytes was also observed in tumors treated with RT and MT1 combination (Fig. 5A). Interestingly, TAMs and, to a lesser extent, Ly6C^high^ monocytes increased their cell membrane levels of PD-L1, likely reflecting their increased activation states (Fig. 5B).

**Figure 5:**
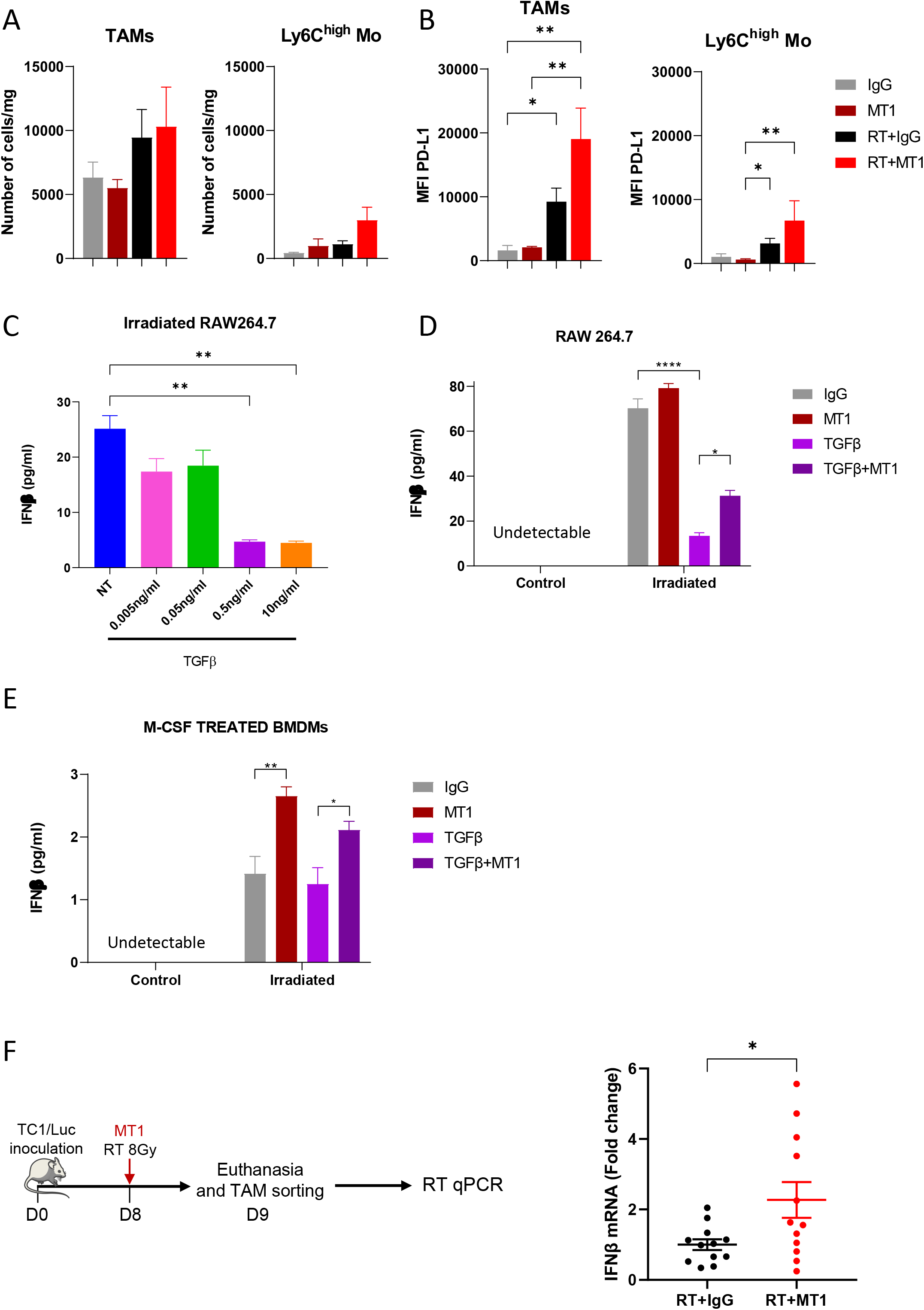
TGFβR2 inhibition triggers the production of IFNβ by macrophages following irradiation. (**A**) Histograms representing the number of TAMs and Ly6C^high^ monocytes/mg of TC1/Luc tumor. (**B**) Mean fluorescence intensity (MFI) of TAMs and Ly6C^high^ monocytes expressing PD-L1 in the different indicated groups (for A and B: n = 8-9 mice from 3 independent experiments, mean ± SEM is reported). (**C**) Histogram representing the production of IFNβ by irradiated RAW264.7 cells determined by ELISA after treatment with different doses of TGFβ. (**D**) Production of IFNβ by RAW264.7 cells with or without irradiation and treated or not with 0.5 ng/ml TGFβ and/or MT1 (for C and D, data from 2 independent experiments performed in duplicate, mean ± SEM is reported). (**E**) Quantification of IFNβ production by BMDMs differentiated with M-CSF for 5 days and then irradiated or not and treated with 0.5 ng/ml TGFβ and/or MT1 (n=8-11 from 3 independent experiments). (**F**) Mice bearing TC1/Luc tumors were irradiated and treated with or without MT1 before euthanasia and tumor macrophages sorting (left panel). The right panel represents the quantification of IFNβ RNA in the extracted macrophages analyzed by qPCR. For all panels: *: p<0.05; **: p<0.01; ****: p<0.0001 (A,B Kruskal-Wallis with Dunn’s multiple comparison test; C,D, E one-way ANOVA with Šidák’s multiple comparison test, F Welch’s t test).

As RT is known to trigger the STING-Type I IFN pathway, we verified whether TGFβ could modulate IFNβ secretion by myeloid cells. As shown in Fig. 5C, treatment of the macrophage cell line RAW264.7 with TGFβ at 0.5 ng/ml or higher significantly decreased the concentration of IFNβ in the supernatant of *in vitro* irradiated cells. Incubation with MT1 was sufficient to partially restore IFNβ production by irradiated RAW264.7 cells, with the IFNβ concentration significantly increased in cells treated with TGFβ and MT1 compared to TGFβ alone (Fig. 5D). We next differentiated bone marrow-derived macrophages (BMDMs) towards an immunosuppressive phenotype using high concentration of M-CSF [25]. Supernatant of irradiated BMDMs contained low amounts of IFNβ that were not reduced further by TGFβ treatment (Fig. 5E). Of note, MT1 treatment increased the secretion of IFNβ by the irradiated BMDMs (Fig. 5E), indicating that TGFβR2 inhibition can trigger the production of IFNβ by immunosuppressive macrophages. Importantly, qPCR analysis of mRNA extracted from TAMs sorted from TC1/Luc oral tumors one day after treatment showed that the inhibition of the TGFβ pathway mediated by MT1 treatment triggered the upregulation of IFNβ transcripts post-RT (Fig. 5F). We also confirmed the significant increase of IFNβ mRNA in macrophages sorted from SCC VII head and neck and LL2/Luc lung tumors treated by the combination of RT and MT1 (Supp. Fig. 4A-B). These data demonstrate that TGFβ in the tumor environment impairs the type I IFN response mediated by irradiated macrophages, and that treatment with MT1 can restore their IFNβ production.

### Type I interferon pathway mediates immune-stimulating antitumor effects of TGFβR2 inhibition

To functionally assign a role to IFNβ in the response to RT and TGFβR2 inhibition, we treated TC1/Luc tumor-bearing mice with anti-type I IFN receptor subunit 1 neutralizing antibody (anti-IFNAR). While anti-IFNAR treatment slightly affected the tumor growth and survival of mice treated with RT alone, the efficacy of treatment combined with RT and MT1 was severely impaired (Supp. Fig. 5A). The survival benefit mediated by MT1 in irradiated mice was completely abrogated when treated with anti-IFNAR (Fig. 6A), indicating that a functional type I interferon pathway is essential to convey the effects of TGFβR2 inhibition in irradiated mice. Accordingly, the viable tumor area as quantified by a head and neck pathologist on histological sections was not reduced when anti-IFNAR was added to the combination of RT and MT1 (Fig. 6B). Additionally, there was no increase in vessel area (Supp. Fig. 5B) and the reduction in tumor hypoxia was limited (Supp. Fig. 5C). The loss of efficacy of MT1 in mice treated with anti-IFNAR was accompanied by severely reduced tumor infiltration by CD8^+^ T cells (Fig. 6C-F), including within the tumor core (Fig. 6F), indicating that the abrogation of TGFβ-mediated immune exclusion upon TGFβR2 inhibition is mediated by the activity of type I interferon. Importantly, we also confirmed the pivotal role of type I IFN pathway in mediating the effects of RT and MT1 combination in the SCC VII head and neck (Supp. Fig. 5D) and in the LL2/Luc lung (Supp. Fig. 5E) tumor models.

**Figure 6:**
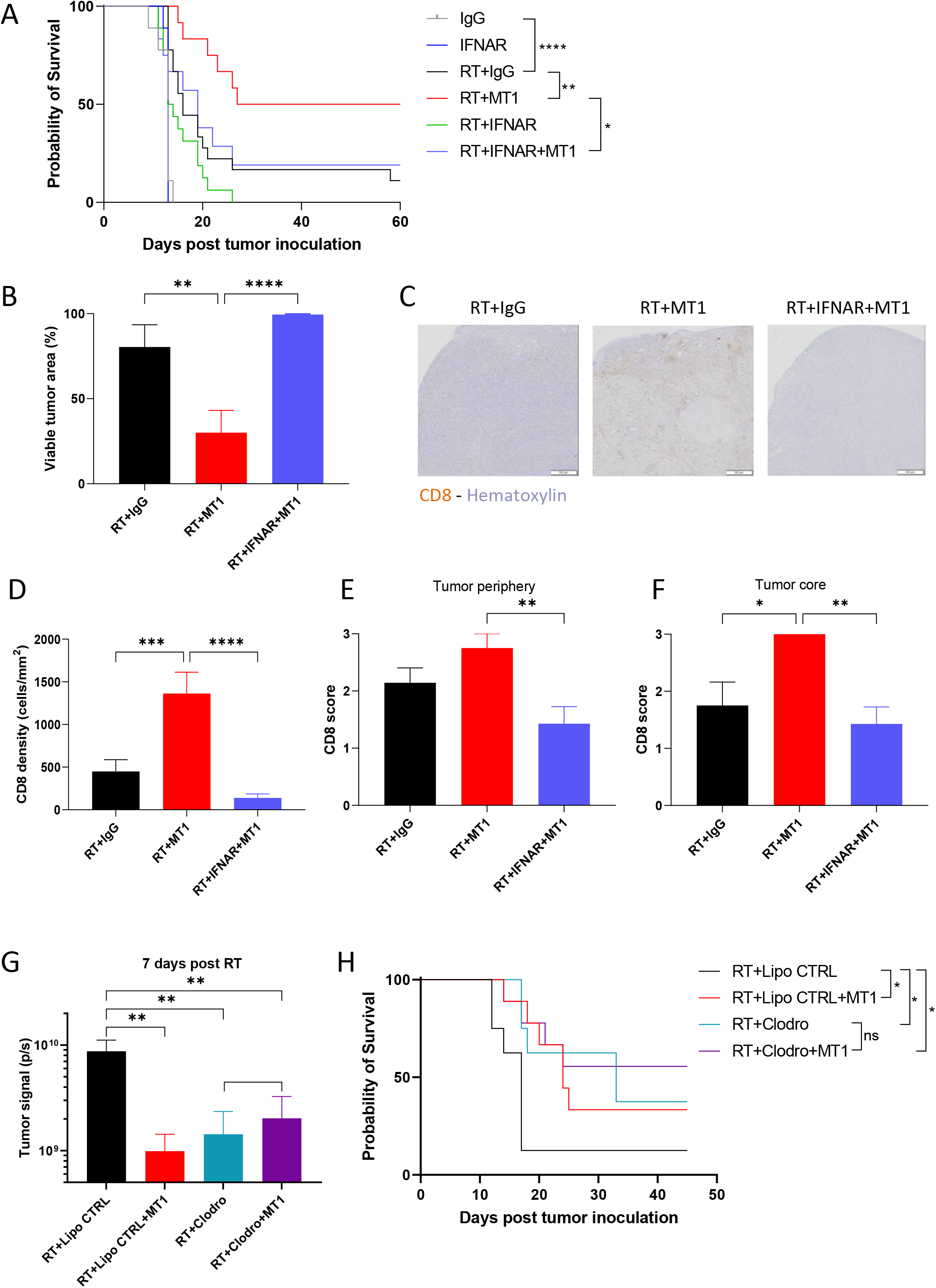
The type I interferon pathway mediates the immune-stimulating antitumor effects of TGFβR2 inhibition. (**A**) Kaplan-Meier mouse survival curves of the different TC1/Luc groups treated with anti-type I IFN receptor subunit 1 antibody (IFNAR) or isotype control (IgG, n = 6-12 mice from 2 independent experiments). (**B**) Histogram representing the percent of viable tumor for each group. (**C)** Representative images of CD8 staining by immunochemistry on head and neck tumors for the RT+IgG (left), RT+MT1 (middle) and RT+IFNAR (right) groups. (**D**) Quantification of total CD8^+^ T cell density per mm^2^ in head and neck tumors. (**E-F**). Semiquantitative scoring of the CD8 density at the tumor periphery (E) and at the tumor core (F) (For B, D-F, n=7-9 from 2 independent experiments). (**G**) Tumor signals were quantified by bioluminescence *in vivo* imaging 7 days post radiotherapy for the different groups of irradiated mice that received intratumor clodronate liposomes (Clodro) or the control PBS liposomes (Lipo CTRL) and/or MT1. (**H**) Kaplan-Meier survival curves of the different groups of irradiated mice that received intratumor clodronate liposomes or the liposome controls and/or MT1 (For G-H, n=8-9 mice from 2 independent experiments). For all panels: *: p<0.05; **: p<0.01; ***: p<0.001; ****: p<0.0001 (A,H log-rank test, B and D-G one-way ANOVA with Tukey’s multiple comparison test).

Finally, we determined the functional role of tumor macrophages in the response to RT and MT1 by depleting them in irradiated TC1/Luc tumors *via* intratumor injection of clodronate liposomes (clodrosomes, Fig. 6G-H). TAM depletion by clodrosomes was sufficient to improve the response to RT when compared to mice treated with PBS-loaded control liposomes (Fig. 6G-H), in agreement with previous reports [22]. The beneficial effect of MT1 in combination with RT was lost in clodrosomes-treated mice, with no significant reduction of tumor size (Fig. 6G) nor improvement of mouse survival (Fig. 6H) in RT+clodrosomes+MT1 mice when compared to RT+clodrosomes mice, in contrast with the significant improvement observed in RT+control liposomes+MT1 mice when compared to RT+control liposomes mice, as expected. These data assign a functional role to TAMs in mediating the effects of the combination of RT and MT1.

## DISCUSSION

A main factor for effective anticancer therapy is its ability to induce an immunogenic antitumor response. Several preclinical data have shown that RT can promote an antitumor immune response, and it has been proposed to behave as an *in situ* vaccine [27]. Nevertheless, highly heterogeneous changes in the tumor immune landscape were reported in cervical cancer patients during the course of concurrent chemoradiotherapy (CCRT), demonstrating that immune activation after CCRT occurs only in a subset of patients, while others even experience a weakening of immune markers during treatment [28,29]. Accordingly, several mechanisms can limit RT efficacy: in addition to non-inflamed phenotypes (cold tumors), immune-excluded and immunosuppressed environment (strongly relying on TGFβ) contributes to therapeutic failure [30,31]. Reversion of immunosuppression in the tumor microenvironment is a key step for successful anticancer therapy. TGFβ has been suggested as a central player in the contribution to tumor resistance [5,14], limiting the efficacy of either radio/chemotherapies [32] or immunomodulators [33]. Here, we showed that the TGFβ pathway represents a pivotal factor of resistance to radiotherapy in two murine models of oral cancer and in an orthotopic model of lung cancer. Employing clinically available monoclonal antibodies and small molecule inhibitors of TGFβ receptors, we demonstrated that targeting the TGFβ pathway improved the efficacy of RT, in line with similar observation in breast [12,34], brain [35] and colorectal [9,34,36] cancer models, and identified a novel mechanism that implies the inhibition of IFNβ production by irradiated macrophages by TGFβ. Of interest, the effect of TGFβ inhibition in combination with RT was observed in both HPV-transformed (TC1/Luc) and HPV-negative (SCC VII and LL2/Luc) models. Our data were obtained using the RT single doses of 8Gy or 12Gy. Since the immunomodulatory effects of radiotherapy have been shown to be dependent on the dose, as well as on the fractionation scheme [37,38], it will be of interest to evaluate if other RT doses/fractions modify the response to the MT1. We also demonstrated, using blocking antibodies against the interferon-α/β receptor (IFNAR), that the improved efficacy of RT observed in TGFβ-treated mice was type I interferon dependent in the TC1/Luc and SCC VII head and neck and in the LL2/Luc lung orthotopic models.

Type I interferon plays a major role in the tumor response to therapies, as it contributes to the induction of a powerful adaptive immune response. After RT, DNA damage followed by dsDNA accumulation in the cytosol triggers the DNA sensing pathway via cGAS/STING, which stimulates the secretion of IFNβ [39]. Here, we showed that *in vitro* irradiation of a macrophage-like cell line is sufficient to trigger significant IFNβ production, which is in agreement with previous reports [40], and that this process is negatively regulated by the TGFβ pathway. Since it has been shown that TGFβ can limit IFNβ production by macrophages stimulated with a STING agonist through the inhibition of IRF3 phosphorylation [41], the activity of TGFβ on irradiated macrophages likely occurs in a similar way. We also showed that in primary BMDMs differentiated towards an anti-inflammatory phenotype the production of IFNβ was low even after RT (when compared to what observed with the RAW264.7 cells) and was not further lowered by treatment with recombinant TGFβ, suggesting that this pathway was already activated in these immunosuppressive macrophages. Accordingly, the impaired IFNβ secretion in M-CSF-differentiated BMDMs and in TGFβ-treated RAW264.7 cells could be effectively reverted using the anti-TGFβR2 antibody MT1, which also upregulated IFNβ expression in TAMs sorted from irradiated tumors. Of note, the low but significant increase of the IFNβ transcript in TAMs from mice treated with MT1 was consistently observed in all three tumors models. Together with the observation that the effect of MT1 was abrogated by the depletion of tumor macrophage, these data strongly support the hypothesis that IFNβ-producing macrophages play a significant role in the tumor response to therapy.

Type I IFNs can act as a link between innate and adaptive immune responses, regulating the capacity of DCs to prime CD8^+^ T cells and generating tumor antigen-specific CD8^+^ T cells [42]. In agreement with the increased activation of IFNβ following treatment with RT+MT1, we observed strong CD8^+^ T cell infiltration, which was type I interferon-dependent, as it was completely abrogated following anti-IFNAR administration. These tumor-infiltrating CD8^+^ T cells were required to mediate the effect of the combination of RT and MT1, which was lost after anti-CD8 antibody treatment. Of note, the combination therapy turns the poorly infiltrated tumors into “hot” tumors through the recruitment of CD8^+^ T cells throughout all tumor tissue, including the core of the tumors, while it was restricted mostly to the tumor periphery for the mice treated with RT alone. Accordingly, differential densities of CD8^+^ T cells in the tumor core and at the invasive margin had prognostic value for survival and response to chemotherapy in breast cancer patients [43,44]. Histological analysis of the spatial infiltration patterns in different cancer types, including HNSCC, showed a heterogeneous “topography” of immune cells with a proportion of immune-excluded tumors [45], thus underlying the importance of modulating immune infiltration to optimize cancer treatment. It was shown in subcutaneous tumor settings that TGFβRI inhibition can strongly promote tumor infiltration by CD8^+^ T cells, through a mechanism involving the upregulation of their CXCR3 levels, which allows for improved homing to CXCR3 ligands produced in the tumor microenvironment following RT [9], and we may speculate that a similar mechanism occurs in our models. In addition, we also observed a trend for increase of the area of tumor vasculature, associated with a reduction of the hypoxia, which might suggest that MT1 increases the tumor accessibility to immune cells, thus likely contributing to the induction of the complete responses observed in the combination group. It has been shown that macrophages can contribute to the T cell exclusion from tumors: in human lung tumors, macrophages mediated lymphocyte trapping by forming long-lasting interactions with CD8 T cells, and in mouse models they limited CD8 T cell migration and infiltration into tumor islets [46]. These data are in line with our observations, as it is thus conceivable to speculate that affecting the macrophage phenotype by TGFβR inhibition could have an impact on the macrophages-CD8 T cell interaction resulting in an improved tumor infiltration.

Together with an increased infiltration of CD8^+^ T cells in mice treated with the combination of RT and MT1, we also observed an increase in tumor NK cells. In agreement, TGFβ is known to limit the membrane expression of the chemokine receptors CXCR3, CXCR4 and CX3CR1, thus affecting NK cell migration and recruitment [47]. As TGFβ is also a powerful negative regulator of NK cell function [48], TGFβR2 inhibition could restore the activity of tumor-infiltrating NK cells, which could contribute to the antitumor efficacy of the combined treatment. On the other hand, we also observed a trend of an increase in the level of Tregs in tumors irradiated and treated with MT1, even if TGFβ is known to promote Treg differentiation in tumors [49]. This observation is in line with a recent report from De Martino et al. [50], who observed an increase of Tregs in subcutaneous tumors treated with RT and TGFβ blockade, which was mediated by upregulation of the TGFβ family member activin A. Thus, HNSCC and lung carcinomas could likely benefit from a TGFβ/activin A double blockade to prevent Tregs accumulation.

Our data demonstrate that the TGFβ pathway acts as a regulator of the radiation-induced adaptive immune response, impairing the synthesis of IFNβ by TAMs and preventing the consequent type-I interferon-driven massive infiltration of the tumor by activated CD8^+^ T cells. Given the several immunomodulatory effects observed in our models, and the induction of an immune-memory following the MT1 and RT combination treatment, it will be of interest in future studies to evaluate if TGFβR inhibition with radiotherapy can also trigger the antitumor response in out of field (abscopal) tumors, as it was previously shown with other molecules impacting the TGFβ pathway [12,34]. These data reinforce the notion that combined immunotherapies are needed to overcome tumor immune suppression and to optimize the efficacy of the treatment.

Overall, our data contribute to elucidating the role of TGFβ in limiting the efficacy of radiotherapy by demonstrating that TGFβ-mediated inhibition of macrophage-derived type I interferon production can impair the adaptive immune response in irradiated tumors.

## Declarations

### Ethics approval

Animal procedures were performed according to protocols approved by the Ethical Committee CEEA 26 and the Ministère de l’Enseignement Supérieur et de la Recherche (MESR) and in accordance with recommendations for the proper use and care of laboratory animals.

### Author contributions

P.H. performed experiments, analyzed results and wrote the manuscript. M. G-d-T. performed experiments, analyzed results and revised the manuscript. M.C. analyzed results and revised the manuscript. N.S. analyzed results. W.L. performed experiments. L.M., C.C. and F.M. provided technical support and revised the manuscript. E.D. designed and supervised the study and revised the manuscript. M.M. designed and supervised the study, performed experiments, analyzed the results and wrote the manuscript.

## Acknowledgments

The authors thank Hélène Rocheteau (PETRA platform), Yann Lecluse, Philippe Rameau and Cyril Catelain (PFIC platform), Patrick Gonin and Karine Ser-Le-roux (PFEP platform) at Gustave Roussy for technical assistance, C. Brunner (Universitätsklinik Ulm, Ulm, Germany) for providing the SCC VII cells and Nadège Bercovici for the fruitful discussions.

## Author information

P.H.’s current address is Department of Oncological Sciences, The Immunology Institute, Tisch Cancer Institute, Icahn School of Medicine at Mount Sinai, New York, NY, 10029, USA.

## List of Abbreviations

TGFβ: transforming growth factor-beta
RT: radiotherapy
TGFβR2: TGFβ receptor
HNSCC: head and neck squamous cell carcinoma
IFN: interferon
TAMs: tumor-associated macrophages
Tregs: regulatory T cells
EMT: epithelial-to-mesenchymal transition
TME: tumor microenvironment
CTL: cytotoxic CD8 T cell
LAP: latency-associated peptide
ROS: reactive oxygen species
IFNAR: interferon-α/β receptor
BMDMs: bone marrow derived macrophages
CCRT: chemoradiotherapy
Tconv: conventional T cells
NK: natural killer cells
DCs: dendritic cell
MFI: mean fluorescence intensity

