## Supplementary material for "TGFβ receptor inhibition unleashes interferon-β production by tumor-associated macrophages and enhances radiotherapy efficacy": Supp. Fig.

**Supplementary Figure 1: TGFβR1 inhibition improves radiotherapy efficacy against head and neck orthotopic tumors.** (**A**) Quantification of the bioluminescent signal from individual tumors was performed at different time points post treatment in mice treated with the TGFβR1 inhibitor LY3200882, RT or RT+ LY3200882 7 days after TC1/Luc inoculation (n = 12-19 mice/group from 2 independent experiments). (**B**) Kaplan-Meier survival curves. (**C**) Tumor signals were quantified by bioluminescence *in vivo* imaging 7 days post radiotherapy for each indicated group. For all panels: *: $p<0.05$; ***: $p<0.001$; ****: $p<0.0001$; ns: non-significant (**B** log-rank test, **C** one-way ANOVA with Tukey's multiple comparison test).

**Supplementary Figure 2: Combination of RT and MT1 modulates tumor vascularization and hypoxia.** (**A**) Histogram representing the percent of TC1/Luc tumor vascularization, representing the % of the area covered by CD31-positive vessels by IHC for each group. (**B**) Representative images of the co-staining of CD8 (pink) and CD31 (red) by IHC on head and neck tumors. (**C**) Histogram shows the density of CAIX+ cells by IHC to quantify hypoxia in the tumors of the different groups. (**D**) Representative images of the CAIX staining. For A-B, D, n=8-9 mice from 2 independent experiments, **: $p<0.01$; ***: $p<0.001$ (Kruskal-Wallis with Dunn's multiple comparison test).

**Supplementary Figure 3: Immunodeficient mice do not respond to the RT+MT1 combination.** (**A**) Quantification of the bioluminescent signal from individual tumors was performed at different time points post treatment in immunodeficient nude mice. (**B**) Kaplan-Meier mouse survival curves. For all panels: n = 7-8 mice/group from one experiment, ***: $p<0.001$; ****: $p<0.0001$.

**Supplementary Figure 4: TGFβR2 blockade induce IFNβ production by macrophages following radiotherapy in head and neck and lung tumor models.** (**A**) Mice were injected with SCC VII cells at a submucoal site in the inner lip to establish a head and neck model. They were then irradiated and treated with or without MT1 before sacrifice and tumor macrophages sorting (left panel). The right panel represents the quantification of IFNβ mRNA in the sorted tumor macrophages analyzed by qPCR. (**B**) Orthotopic lung tumor model was established by transpleural injection of LL2/Luc in the lung. Mice were then irradiated and treated or not with MT1 before sacrifice and macrophage sorting (left panel). The right panel shows the quantification of IFNβ mRNA in the sorted tumor macrophages analyzed by qPCR. For all panels: **: $p<0.01$; (Welch's t test).

**Supplementary Figure 5: Anti-IFNAR administration impairs the RT and MT1 combination efficacy.** (**A**) Quantification of the bioluminescent signal from individual TC1/Luc tumors was performed at different time points post treatment in the different groups treated with anti-IFNAR (IFNAR) antibody or isotype control (IgG) (n = 6-12 mice from 2 independent experiments). (**B**) Quantification of the tumor vascularization (% of the area covered by CD31-positive vessels) by IHC for each group. (**C**) Histogram shows the density of CAIX positive cells by IHC to quantify hypoxia in the tumors of the different groups. (**D**) Kaplan-Meier survival curves of SCC VII head and neck model in the different groups (n=5-12 mice per groups from 2 independent experiments). (**E**) Kaplan-Meier survival curves of LL2/Luc orthotopic lung tumor model in the different groups (n=4-6 mice per groups from 2 independent experiments). For all panels, *: $p<0.05$; **: $p<0.01$; ***: $p<0.001$; ****: $p<0.0001$ (C one-way ANOVA with Tukey's multiple comparison test, D-E log-rank test).

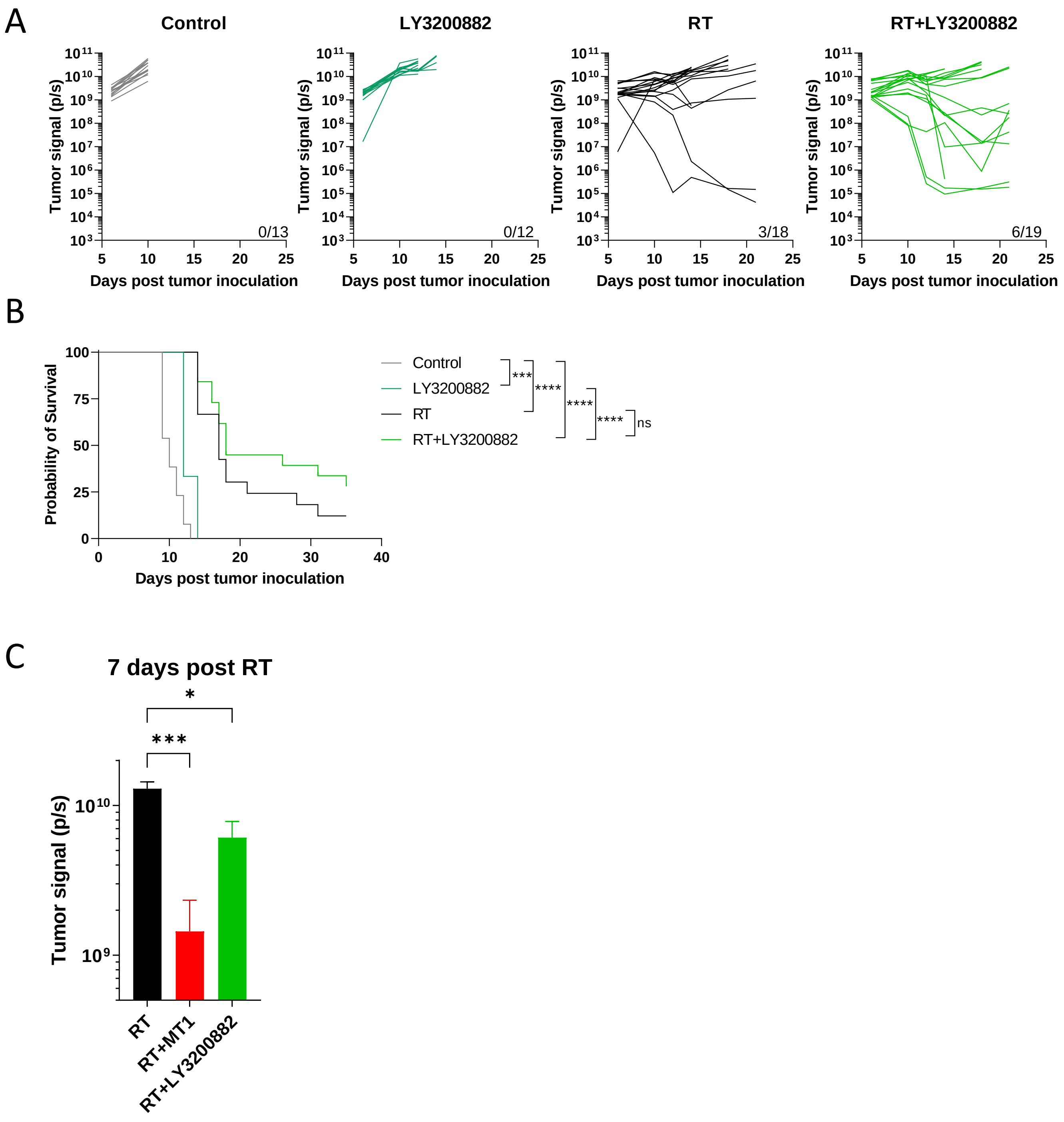


Supplementary Figure 1

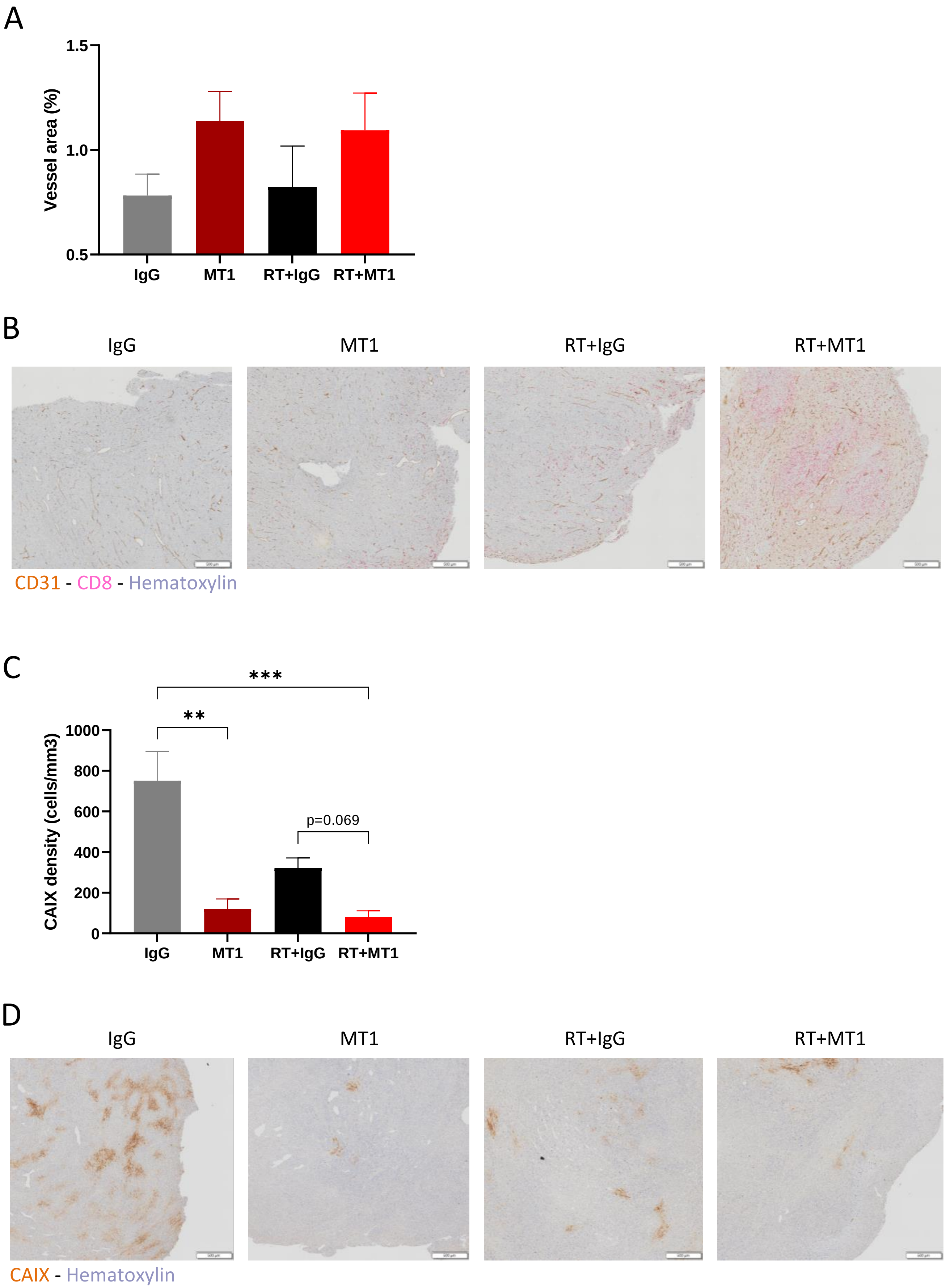


Supplementary Figure 2

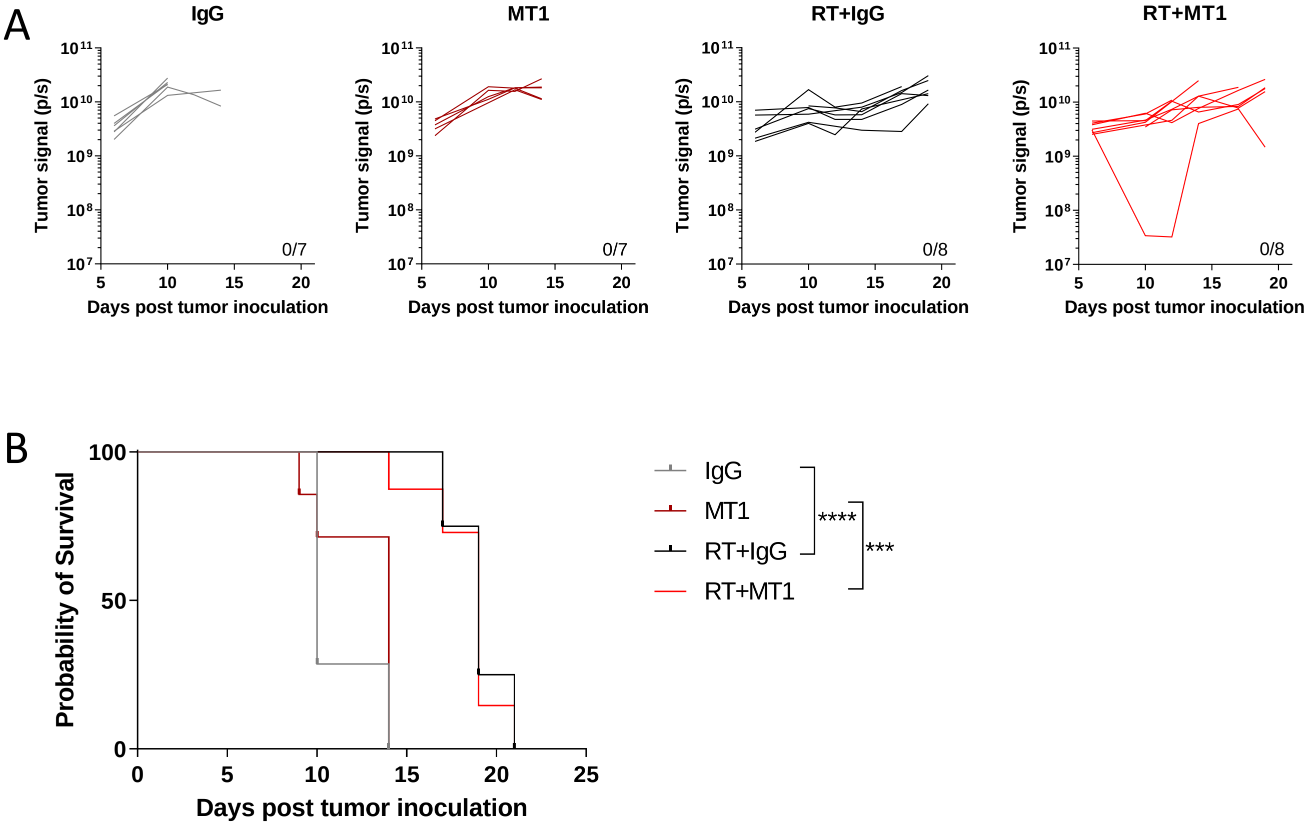


Supplementary Figure 3

A

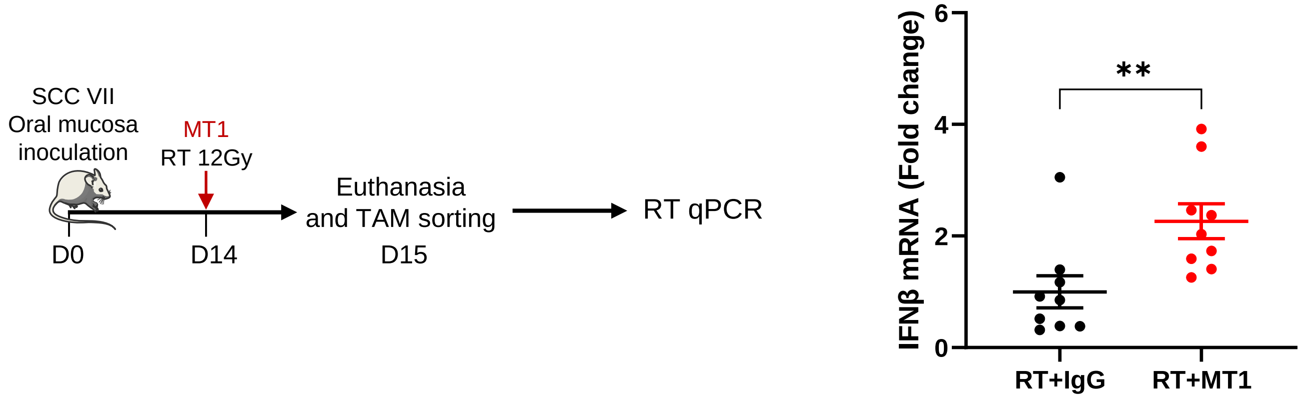


B

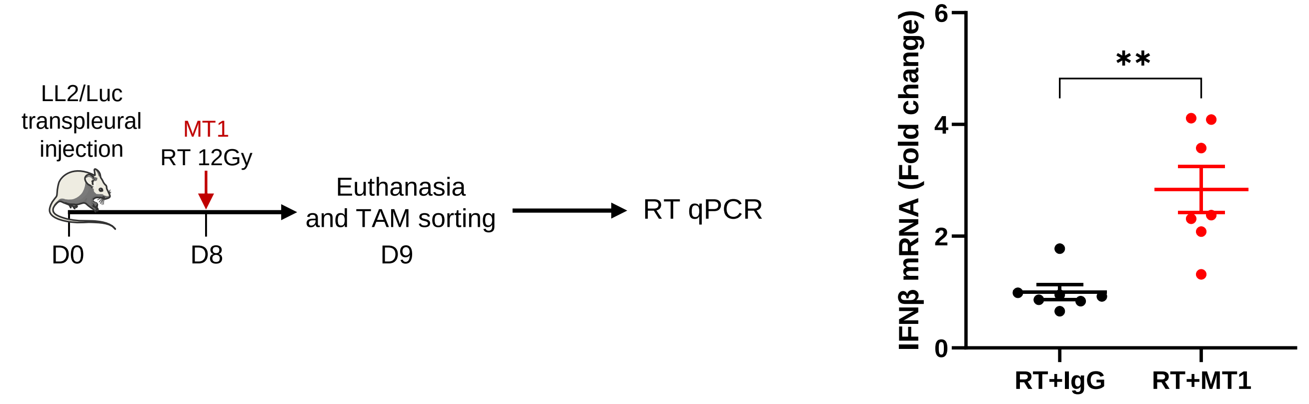


Supplementary Figure 4

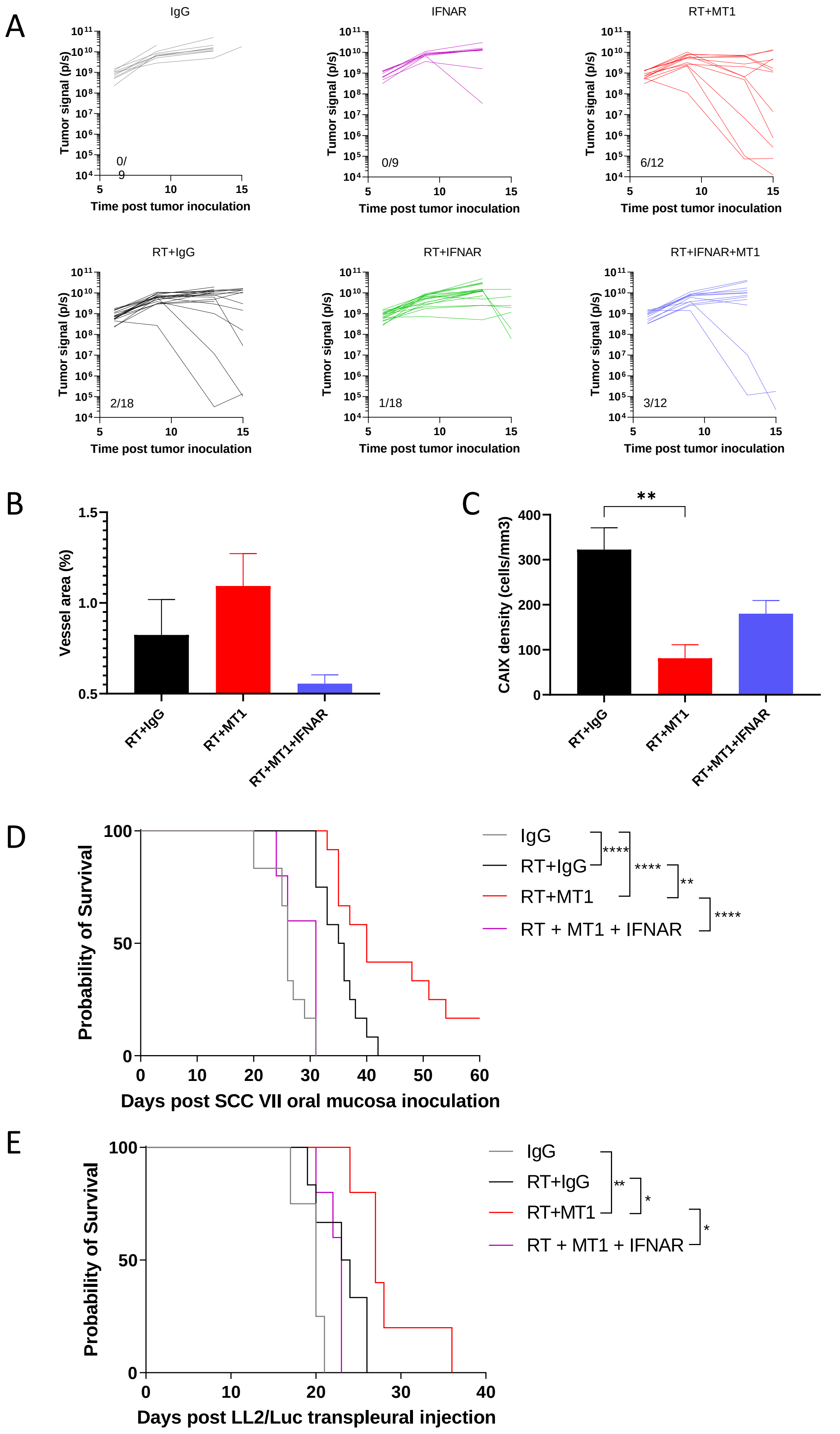


Supplementary Figure 5
